# Type 6 secretion systems of adherent-invasive *Escherichia coli* isolated from Crohn’s disease patients shape intestinal microbiota

**DOI:** 10.64898/2026.10.08.757420

**Authors:** Hélène Botchorichvili, Sébastien Massier, Benoit Chassaing, Aurélien Birer, Caroline Chevarin, Gwladys Sevrin, Michael Rodrigues, Sylvain Denis, Ivan Wawrzyniak, Eric Peyretaillade, Richard Bonnet, Bruno Pereira, Matthieu Allez, Philippe Seksik, Jérémy Denizot, The Remind Group, Anthony Buisson, Laure Journet, Nicolas Barnich, Elisabeth Billard

**Author notes:** Corresponding author : Elisabeth BILLARD, UMR1071 Inserm M2iSH, 28 place Henri Dunant 63000 Clermont-Ferrand France. These authors contributed equally to this work (Hélène Botchorichvili, Sébastien Massier, Nicolas Barnich, Elisabeth Billard). PBS (Polymers, Biopolymers, Surfaces), CNRS UMR 6270, Université de Rouen Normandie, INSA de Rouen Normandie, Rouen, France. **Dedication:** We wish to dedicate this work to our colleague and friend Nicolas Barnich, who sadly passed away before its publication. Nicolas was a brilliant researcher and dedicated mentor whose contributions to the AIEC field and to this work were decisive. He mentored several of us, and there are no words to express how deeply we miss him. Remind Study Group investigators: Christophe Attencourt, Clotilde Baudry, Laurent Beaugerie, Madeleine Bezault, Gilles Bommelaer, Gilles Boschetti, Yoram Bouhnik, Anne Bourrier, Sarah Boyer, Franck Brazier, Loic Brot, Marine Camus, Pierre Cattan, Najim Chafai, Ulriikka Chaput, Anne-Laure Charlois, Denis Chatelain, Marion Chauvenet, Mircea Chirica, Dilek Coban, Olivier Corcos, Hélène Corte, Eddy Cotte, Clotilde Debove, Pierre Desreumaux, Marie Dodel, Xavier Dray, Anne Dubois, Jean-Louis Dupas, Peggy Falgon, Mariane Ferron, Jérome Filippi, Bernard Flourié, Yves François, Mathurin Fumery, Jean-Marc Gornet, Marion Goutte, Nathalie Guedj, Nassim Hammoudi, Xavier Hébuterne, Paul Hofman, Nadia Hoyeau-Idrissi, Julien Kirchgesner, Cecilia Landman, Martine Leconte, Jeremie Lefevre, Nelson Lourenco, Leon Maggiori, Marianne Maillet, Carole Margalef, Philippe Marteau, Driffa Moussata, Nicolas Munoz-Bongrand, Maria Nachury, Stéphane Nancey, Isabelle Nion-Larmurier, Andrée Nisard, Yves Panis, Yann Parc, Charles Sabbagh, Harry Sokol, Coralie Sommeville, Carmen Stefanescu, Virginie Suffys, Magali Svrcek, My-Linh Tran-Minh, Xavier Treton, Pauline Wils, Clara Yzet, Philippe Zerbib.

## Abstract

Adherent Invasive *Escherichia coli* (AIEC) colonize the ileal mucosa of Crohn’s disease patients. Among bacterial virulence factors potentially implicated, type 6 secretion systems (T6SS) are molecular machineries sending effectors in prokaryotic or eukaryotic target cells. Two T6SS have been reported in AIEC LF82 reference strain. Here, we show that T6SS genes are more prevalent in Crohn’s disease patients’ microbiota compared to healthy controls, and that T6SS are more frequent in AIEC pathovar compared to *E. coli* in general and associated with FimH pathoadaptive polymorphism. The use of an *in vitro* gastrointestinal system coupled with T6SS promoter reporter assay highlighted expression of LF82 T6SS under gastrointestinal conditions, with a major role for bile salts. In mice infected with AIEC LF82 *versus* T6SS-deleted mutants, T6SS showed a functional impact in modulating the composition of mucosa-associated gut microbiota. Thus, AIEC may use their T6SS to compete with endogenous microbiota to colonize host mucosa.

## Introduction

Crohn’s disease (CD), one of the inflammatory bowel diseases, is a chronic condition that develops progressively and evolves in a relapsing and remitting manner, causing inflammation and lesions on all segments of the gastrointestinal tract. The most affected sites are the intestinal mucosae in the terminal ileum and the colon. To date, no curative treatments exist and CD patients can only benefit from symptomatic treatment to limit the frequency and the intensity of inflammation flares. Although CD incidence is stabilizing in western countries, it remains a major public health problem, with high prevalence in Europe and North America and rising incidence in newly industrialized countries^1,2^. Etiology of CD is multifactorial, resulting from the interplay between genetic susceptibility, environmental factors and abnormal intestinal microbiota composition.

Indeed, dysbiosis observed in CD patients is characterized by a reduction of global microbial diversity and, more specifically, a decrease in beneficial bacteria (eg, Bacillota phylum, formerly Firmicutes) and an increase of Bacteroidota and Pseudomonadota phyla (formerly Bacteroidetes and Proteobacteria). Enterobacteriaceae, members of the Pseudomonadota phylum, can represent up to 100% of anaerobic ileal mucosa-associated microbiota. Incidentally, numerous studies revealed that some *Escherichia coli* strains named Adherent Invasive *E. coli* (AIEC) were significantly more abundant in CD patients and mainly associated with the inflamed ileal mucosa. Indeed, the prevalence of AIEC in the mucosa of adult patients with CD ranges from 21% to 63% versus 6% to 19% in healthy subjects^3,4^. Those strains have two main characteristics, the capacity to adhere/invade epithelial cells and the ability to survive in macrophages^5,6^.

Genome analysis of the AIEC reference strain LF82 revealed the presence of pathogenic islands encoding two type six secretion systems (T6SS)^7,8^. T6SS is a multiprotein nanomachinery that can deliver effectors into target cells using a spike-like mechanism. This injection apparatus, analogous to T4 bacteriophage tail-like structure, is composed of a membrane complex, a baseplate, and a tail tube/sheath complex topped by a molecular spike^9^. When the sheath contracts, the tube topped by the spike is propelled outside, possibly puncturing the membrane of other cells either eukaryotic or prokaryotic. Perforation enables effectors loaded on the system to be delivered directly into the target cell. First identified in *Vibrio cholerae*, T6SS are usually encoded within a single genetic locus on the chromosome composed of a minimum of 13 genes necessary to assemble a functioning apparatus^10–12^, to which is added a Proline-Alanine-Alanine-aRginine (PAAR)-coding gene. PAAR proteins are essential components of the spike and required for T6SS function^13,14^. Those secretion systems are largely common, found in around 25% in Gram-negative Proteobacteria, with an over-representation in *Gammaproteobacteria*^15^.

By phylogenetic analyses, T6SS were classified in four main types according to gene organization and to sequence homologies: T6SS^i^, T6SS^ii^, T6SS^iii^ and T6SS^iv^, among which *E. coli* are reported to carry only T6SS^i^ that have been assigned to subclasses T6SS^i1^ (also called T6SS-2), T6SS^i2^ (also called T6SS-1) and T6SS^i4b^ (also called T6SS-3)^16^. Gene clusters encoding T6SS-1 and T6SS-2 have been previously identified in LF82 genome^7,8^, however their roles in bacterial virulence remain elusive.

This study aimed to extend the present knowledge on T6SS expression conditions in LF82 and on the impact of those systems on intestinal microbiota composition. After measuring expression of both T6SS under intestinal conditions using several *in vitro* models, we identified bile salts as a major activator of the expression of the two systems. Functional impact of LF82 T6SS on mouse intestinal microbiota was assessed *in vivo* in mice by comparing the impact of oral infection with LF82 WT strain or with its isogenic mutant deleted for one (LF82 ΔT6SS1, LF82 ΔT6SS2) or both (LF82 ΔΔT6SS1.2) T6SS-coding operons. At this time, only one T6SS effector had been characterized in LF82 strain : p435, also known as Tle3, carrying a phospholipase A1 activity^17^. The exact function of Tle3 in AIEC virulence remained to be studied, so an isogenic mutant deleted for the gene encoding this effector (LF82 Δ435) was included in our experiments to explore the role of this effector within microbiota.

## Material and Methods

### Bacterial strains, plasmids, and growth conditions

Bacteria and plasmids used in this study are listed in **Table 1**. AIEC strain LF82 is isolated from a chronic ileal lesion of a patient with CD^6^. Bacteria were grown at 30, 37 or 42°C with or without 180 rpm shaking in Luria-Bertani (LB) medium supplemented with the appropriate antibiotic concentration (ampicillin 100 µg/mL, kanamycin 50 µg/mL).

**Table 1.** Strains and plasmids used in this study.

|  | Description <sup>a</sup> |
| --- | --- |
| **Plasmids** |  |
| pST76-K<sup>31</sup> | pSC101ts replication system itself, carries *I-SceI* site (Km<sup>R</sup>) |
| pST76-K-TS1 | pST76 harboring the 945-bp *EcoRI-BamHI* A'B' fragment that frames T6SS1 of LF82 strain |
| pST76-K-TS2 | pST76 harboring the 1274-bp *EcoRI-BamHI* A'B' fragment that frames T6SS2 of LF82 strain |
| pDAI-SceI<sup>32</sup> | pDA17 carrying the *I-SceI* gene (Tet<sup>R</sup>) |
| pRS550 | *E. coli* multicopy lacZ fusion vector (Km<sup>R</sup>, Amp<sup>R</sup>) |
| pRS550-TS1 | pRS550 harboring the 747-bp *BamHI-EcoRI* fragment upstream T6SS1 of LF82 strain |
| pRS550-TS2 left | pRS550 harboring the 747-bp *BamHI-EcoRI* fragment upstream T6SS2 left of LF82 strain |
| pRS550-TS2 right | pRS550 harboring the 1203-bp *BamHI-EcoRI* fragment upstream T6SS2 right of LF82 strain |
| **Strains** |  |
| LF82 WT | *E. coli* LF82 wild-type adherent-invasive (Amp<sup>R</sup>, Ery<sup>R</sup>) |
| LF82 ΔT6SS1 | LF82 mutant with T6SS1 genes deleted |
| LF82 ΔT6SS2 | LF82 mutant with T6SS2 genes deleted |
| LF82 ΔT6SS1.2 | LF82 mutant with T6SS1 and T6SS2 genes deleted |
| LF82 Δ435 | LF82 mutant deleted from gene referenced as 435 |
| LF82-TS1 | LF82 Wild-type with pRS550-TS1 plasmid |
| LF82-TS2 left | LF82 Wild-type with pRS550-TS2 plasmid |
| LF82-TS2 right | LF82 Wild-type with pRS550-TS3 plasmid |

<sup>a</sup> Amp, ampicillin ; Km, kanamycin ; Ery, erythromycin ; <sup>R</sup>, resistance

### Molecular biology

*EcoRI, BamHI, SalI* and *XhoI* endonucleases (New England Biolabs), T4 DNA ligase (New England Biolabs), NucleoSpin®Plasmid kit and NucleoBond® Xtra Midi Kits (Macherey-Nagel) were used according to manufacturer’s instructions. PCR amplifications were performed using Platinium® Taq DNA polymerase High Fidelity (Life Technologies). PCR Master Mix (Euromedex) was used for screening assays and cloning verifications according to the manufacturer’s instructions. Primers used in this study are described in **Table 2**.

**Table 2.**
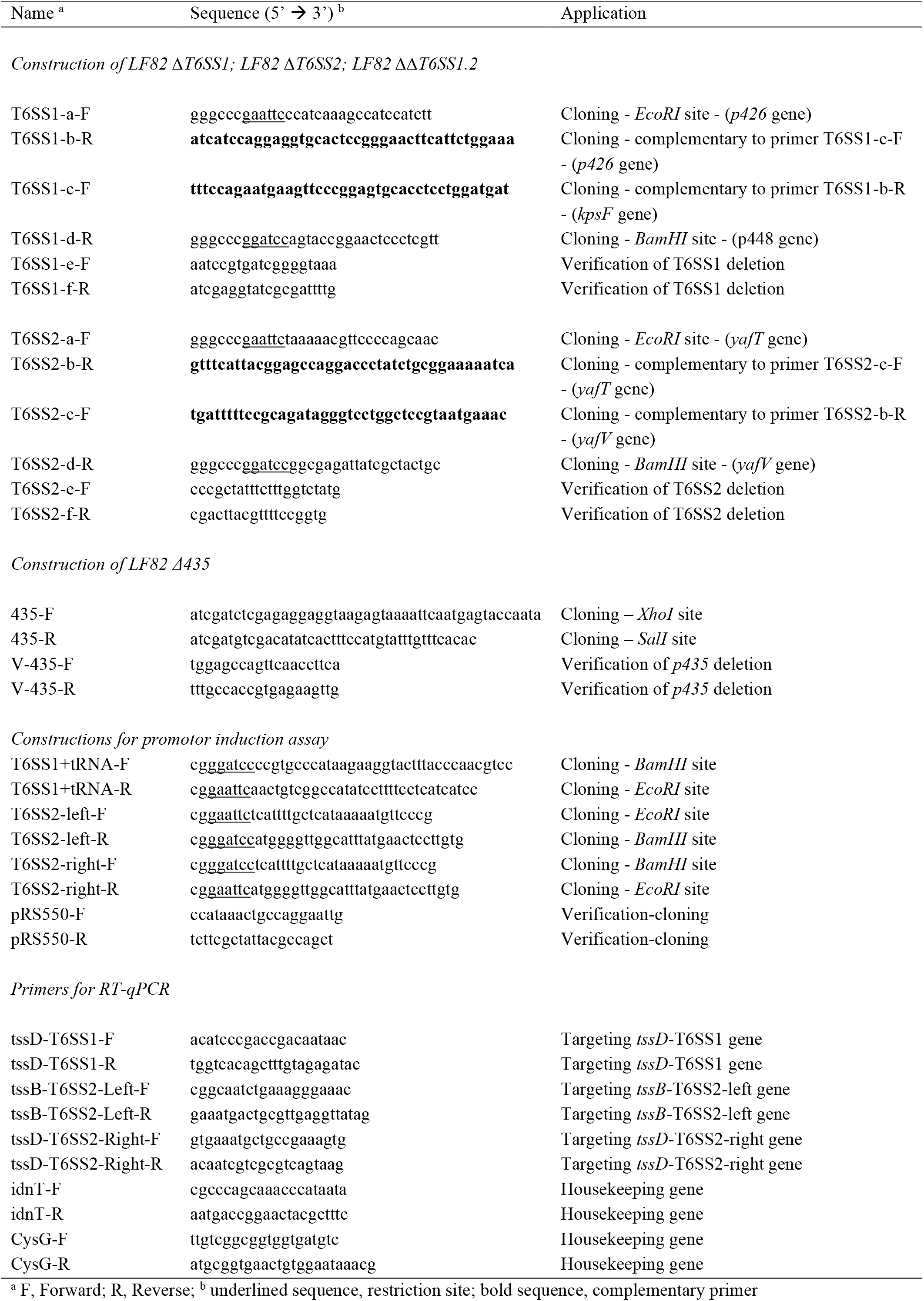
Primers used in this study.

### T6SS prevalence in AIEC

T6SS clusters prevalence was first evaluated on 78 AIEC genomes from our collection (CEALIVE study^18^, POP-Remind study^19^ and Pacific study^20^) with MacSyFinder’s model set TXSScan designed for genomic detection of bacterial secretion systems^21–23^. In a second step, the prevalence of T6SS homologous to those of LF82 was compared between AIEC genomes and 17094 genomes of *E. coli* selected from RefSeq database v1.214k to remove low-quality or redundant sequences. For this purpose, BLASTN analyses were performed using nucleotide sequences with a threshold at 70% coverage and identity, as it allows about 90% probability or more to share the same biological process^24^. Whole clusters were considered when all four genes (*tssB, tssC, tssG* and *tssF* from LF82 genome, accession number: NC_011993) were present with a maximum distance between genes of 34 kb (considering that maximum distance between T6SS genes is 32kb in LF82 genome). Of note, AIEC strains carrying T6SS homologous to that of LF82 have probably been missed in this analysis due to contig discontinuity interrupting the cluster and/or due to homology or coverage lower than 70% with LF82 genes. The prevalence data obtained for T6SS on our AIEC collection were next cross-referenced with those for the S70/N78 polymorphism on FimH adhesin.

### T6SS genes abundance in human microbiomes

Abundance of T6SS genes in human microbiomes of CD patients and healthy controls was assessed using MetaQuery web application^25^ (https://metaquery.docpollard.org/) based on protein sequences encoded by each LF82 T6SS gene with a 80% identity threshold.

### T6SS expression in *in vitro* gastrointestinal tract model TIM-1

The TNO gastro-Intestinal Model (TIM-1) is a dynamic system reproducing the digestive process of human from the stomach to the end of the ileum^26^. Briefly, the main physicochemical parameters of the intestinal environment are reproduced according to physiology and meal conditions: body temperature, pH profiles, intestinal tone, gastric and duodenal secretions (HCl, pepsin, lipase, pancreatic enzymes, bile, NaHCO_3_), gastric and gastrointestinal emptying profiles. A dialysis system is connected to the jejunum and the ileum of the model to mimic passive absorption of water and digested compounds. Here, the TIM-1 has been programmed to simulate the digestive conditions of a healthy adult ingesting a glass of water^27^.

AIEC LF82 bacteria were grown overnight at 37°C without shaking, then 2.10^9^ bacteria were centrifuged, suspended in 200 mL of mineral water, and inoculated into the stomach compartment of TIM-1. The ileal effluents were collected on ice for periods of 1 hour during the 5 hours of the digestion. Total bacterial RNAs were extracted from the inoculum and ileal effluents using a trizol method (Invitrogen) adapted from Toledo-Arana et al.^28^. Genomic DNA was digested using Turbo DNase (Ambion) and then total RNAs were purified using the RNeasy Mini kit (QIAGEN) and quantified with nanophotometer Nanodrop P300 (Implen). RNAs integrity (RNA integrity number-RIN) was checked on Agilent 2100 Bioanalyzer. Next steps of the protocol were performed by Helixio (Saint Beauzire, France). Samples were reverse transcribed using random primers and libraries were then made using the NEB Next Ultra IIRNA Library Prep Kit for Illumina (NEB). Sequencing was performed using an Illumina sequencer (NextSeq500). RNA-seq datasets from 3 independent experiments were randomly down-sampled to a common sequencing depth (total read count: 7 373 080, mean read length = 76 pb) and read counts were normalized for each T6SS gene according to its length.

### T6SS promoter induction assay

Promoter fusion constructs were generated by PCR amplification of T6SS2 left, T6SS2 right and T6SS1-lacZ promoters using T6SS2left F/R, T6SS2-right F/R and T6SS1+tRNA F/R primers respectively (**Table 2**). Resulting fragments containing Shine-Dalgarno sequences as well as translation initiation sites were cloned into plasmid vector pRS550^29^ using *EcoRI*/*BamHI* sites. β-Galactosidase activity was analyzed according to the Miller method based on o-nitrophenyl-β-D galactopyranoside hydrolysis^30^ in strains harboring pRS550-TS1, pRS550-TS2right, pRS550-TS2left or empty pRS550 constructs. Overnight bacterial cultures were resuspended in the following culture media: RPMI supplemented with 10% heat-inactivated FCS, LB medium, LB pH=5 or 9, LB supplemented with NaCl (20 g/L), sucrose (0.5 M), sodium cholate 2% (Sigma Aldrich), bile salts 2% (cholic and deoxycholic acid sodium salt, Sigma Aldrich), polymyxin B (0.25 μM, Sigma Aldrich), 2-2’ bipyridyl (250 μM, Sigma Aldrich) or H_2_O_2_ (1.0 mM, Sigma Aldrich) and in mimic vacuole medium (100 mM bis-Tris, 0.1% casamino Acids, 0.16% glycerol, 10 μM MgCl2, pH adjusted to 5.8 with HCl). ONPG hydrolysis and bacterial concentrations were determined after 3 h by measuring absorbance at 420nm and 620 nm respectively. β-Galactosidase activities were expressed in Miller units^30^.

### RT-qPCR-based quantification of T6SS genes expression

For quantification of structural T6SS genes expression with RT-qPCR, LF82 WT was grown overnight. Bacterial cultures were diluted to OD=1 (600 nm). After centrifugation at 6000 g for 3 minutes, pellets were washed with PBS, recentrifuged and resuspended in LB broth supplemented with 0%, 1% or 2% of bile salts (Sigma-Aldrich) at 37°C for 15 hours without shaking. PBS-washed pellets were dry frozen and kept at -80°C. Bacterial RNA extraction was made with Qiagen® RNeasy kit according to manufacturer’s instructions. Sensifast cDNA synthesis Kit (Bioline) was used for retrotranscription and SsoAdvanced Universal SYBR® Green SuperMix (Bio-rad) was used for qPCR according to manufacturer’s instructions. Primers tssD-T6SS1-F, tssD-T6SS1-R, tssB-T6SS2-Left-F, tssB-T6SS2-Left-R, tssD-T6SS2-Right-F, tssD-T6SS2-Right-R, idnT-F, idnT-R, cysG-F and cysG-R primers are described in **Table 2**.

### Construction of *E. coli* LF82 T6SS mutants

Mutant strains LF82 ΔT6SS1, LF82 ΔT6SS2 and double mutant LF82 ΔΔT6SS1.2 were generated using mutagenesis strategy adapted from Pósfai et al.^31^ and Flannagan et al.^32^ (see **Figure S1** for details and primers in **Table 2**). Briefly, to delete whole T6SS clusters, the upstream and downstream regions of T6SS1 or T6SS2 were amplified by PCR. Then, a fusion PCR was performed to amplify a contiguous fragment. The PCR amplification products were cloned into *EcoRI*/*BamHI* sites of pST76-K which carries a temperature-sensitive replicon, a kanamycin resistance gene (Km^R^) and the recognition site for *I-SceI*. These mutagenesis plasmids (pST76-K-TS1 or TS2) were electroporated into *E. coli* LF82 WT strain, and kanamycin resistant colonies were selected at 30°C. Homologous recombination induced by incubation at 42°C for 5 to 9 h in LB broth allowed plasmid integration into *E. coli* LF82 genome. Upstream or downstream single cross-over insertion of plasmids was verified by PCR. Finally, pDAI-SceI plasmid (gift from M. Valvano, Addgene #32061) carrying tetracyclin resistance gene and constitutively expressing I-SceI nuclease was electroporated into recombinant strains. Loss of the integrated mutagenesis plasmids was verified by picking colonies onto LB agar plates with and without kanamycin. Isogenic mutant LF82 Δ435 was generated by replacement of the p435 sequence with a kanamycin cassette generated by PCR. Deletions were verified by PCR amplification and sequencing.

### *In vivo* mouse experiment

This study was carried out in strict accordance with the recommendations of the Guide for the Care and use of Laboratory Animals of Clermont Auvergne University (Clermont-Ferrand, France). The animal protocol was approved by the Committee for Research and Ethical Issues of the Department of Auvergne (C2EA-02, Auvergne, France; APAFIS #38082). C57BL/6 WT 8-week-old males from Charles River were randomly assigned to experimental groups (n = 6 mice per group). Mice were gavaged for four consecutive days with 200 µL of PBS for the control group or with 10^9^ bacteria in 200 µl of PBS for the infected groups. Body weight was monitored throughout the experiment. Fresh fecal pellets were collected at day 10 and immediately frozen in liquid nitrogen. The data presented are from all the mice included in the experiment (no data excluded) and there was no blinding of the experimenters to the group allocation.

### DNA extraction for microbiota analysis

Bacterial DNA was extracted from frozen fecal samples using the DNeasy 96 PowerSoil Pro QIAcube HT kit (Qiagen), with mechanical disruption via bead beating (TissueLyzer II, Qiagen) as described. Shortly, samples were transferred to 96-well PowerBead plates and mixed with 800 µL of CD1 lysis buffer. Cells were lysed by bead beating for 5 min using the TissueLyzer II to release DNA. Plates were then centrifuged at 2,773 × g for 10 min at 20 °C to pellet debris and beads. A volume of 500 µL of the supernatant was transferred to an S-block (96-well plate), mixed with CD2 solution, and centrifuged again under the same conditions to remove proteins and inhibitors. Next, 550 µL of the resulting supernatants were transferred to a new S-block, and the following steps were carried out on the QIAcube HT platform: 450 µL of C3 binding buffer was added to facilitate DNA binding to QIAamp 96-well silica membrane plates. Bound DNA was washed sequentially with 1 mL of AW1 buffer, 600 µL of AW2 buffer, and 400 µL of 96% ethanol. DNA was eluted in a total volume of 150 µL using a combination of 120 µL of C6 and 30 µL of Top Elute buffer, and stored at −20 °C until further analysis.

### Microbiota analysis by 16S rRNA gene sequencing using Illumina technology

Microbial community composition was assessed by 16S rRNA gene amplicon sequencing using the Illumina MiSeq platform. The V4 region of the 16S rRNA gene was amplified via polymerase chain reaction (PCR) from each sample using a composite forward primer and a reverse primer, each incorporating a unique 12-base barcode designed with the Golay error-correcting scheme to enable sample-specific tagging of PCR products. The forward primer 515F used for amplification was 5’-*AATGATACGGCGACCACCGAGATCTACACGCT*XXXXXXXXXXXX**TATGGTAATT** *GT*GT<u>GYCAGCMGCCGCGGTAA</u>-3’. The italicized segment corresponds to the 5’ Illumina adapter, the 12 ‘X’ characters represent the Golay barcode, the bold portion denotes the primer pad, the italicized and bold sequence serves as the primer linker, and the underlined section is the conserved bacterial primer 515F. Similarly, the reverse primer 806R used was 5’-*CAAGCAGAAGACGGCATACGAGAT***AGTCAGCCAG*CC***<u>GGACTACNVGGGTWTCTAAT</u>-3’. The italicized sequence represents the 3’ reverse complement of the Illumina adapter, the bold segment corresponds to the primer pad, the italicized and bold sequence functions as the primer linker, and the underlined section is the conserved bacterial primer 806R. PCR amplification was performed using 5PRIME HotMasterMix (Quantabio, Beverly, MA, USA) with a final primer concentration of 0.2 μM and a template DNA input ranging from 10 to 100 ng. The thermal cycling conditions included an initial denaturation at 95 °C for 3 min, followed by 30 cycles of 95 °C for 45 s, 50 °C for 60 s, and 72 °C for 90 s on a Bio-Rad thermal cycler. Amplicons were subsequently analyzed via agarose gel electrophoresis. DNA quantification was performed using the Quant-iT PicoGreen dsDNA assay, and equimolar amounts of PCR products were pooled to generate a master DNA library which was further purified using AMPure magnetic beads (Agencourt, Brea, CA, USA) before sequencing (paired-end reads, 2 × 250 bp) on an Illumina MiSeq platform at the Genom’IC facility, Institut Cochin, Paris, France.

### 16S rRNA gene sequence analysis

The 16S rRNA sequences were analyzed using QIIME2 (version 2022). Sequence demultiplexing and quality filtering were performed using the DADA2 algorithm with default QIIME2 parameters to identify and correct Illumina amplicon sequencing errors, generating a table of QIIME2 artifacts. The following DADA2 command was employed to generate a QIIME2 artifact table: “qiime dada2 denoise-paired --i-demultiplexed-seqs demux.qza --p-trim-left-f 0 --p-trim-left-r 0 --p-trunc-len-f 180 --p-trunc-len-r 180 --o-representative-sequences rep-seqs-dada2.qza --o-table table-dada2.qza --o-denoising-stats stats-dada2.qza --p-n-threads 6” Dada2 command. A phylogenetic tree was subsequently constructed using the align-to-tree-mafft-fasttree pipeline to support phylogenetic diversity calculations. Alpha and beta diversity metrics were computed using the core-metrics-phylogenetic workflow, with beta diversity visualized via Principal Coordinates Analysis (PCoA) plots. Taxonomic classification was performed using a 99% identity threshold against the Greengenes2 reference database (version 2022.10), and features were assigned to operational taxonomic units (OTUs) accordingly. We employed the Microbiome Multivariable Associations with Linear Models (MaAsLin2) approach to identify differentially abundant microbial communities: q-values (False Discovery Rate-adjusted p-value) are corrected for the multiple statistical tests by the Benjamini– Hochberg method and reported on the graphics (q-value thresholds for significance : * q<0.25, ** q<0.1, ***q<0.05).

### POP-Remind cohort description

Clinical data of CD patients from the multicentre POP-Remind cohort (Ileal Crohn’s disease and Post-Operative outcome: Prospective cohort study of the REMIND Group, NCT03458195^33^, registration date March 1st, 2018) were used to assess the impact of colonization with T6SS+ AIEC strains on clinical outcomes. The study was approved by the Agence Française de Sécurité Sanitaire et des Produits de Santé (IDRCB 2009-A00205-52) and by the ethics committee of Saint-Louis Hospital (CPP 2009/17). Briefly, adult patients with CD undergoing ileocolonic resection were enrolled after giving full informed consent. Endoscopic evaluation was performed, clinical data were collected and patients’ AIEC colonization was determined from ileal mucosal biopsies as detailed previously by isolating *E. coli* strains and assessing their adhesion and invasion properties in intestinal epithelial cells^19^.

### Statistical analysis

Except for microbiota analysis, statistical analyses were performed as indicated in figure legends with GraphPad Prism software (Boston, USA) or with Stata software (Stata Corp, College Station, USA). A p-value less than 0.05 was considered statistically significant.

## Results

### T6SS genes prevalence in CD and healthy intestinal microbiota

The two clusters of pathogenicity previously identified in AIEC strain LF82 genome encode T6SS1 and T6SS2, with genes encoding T6SS structural proteins as well as known or putative effectors and their immunity-associated proteins. In addition to previously described operons, we identified some other T6SS-associated genes outside of the two main clusters, in the VgrG island (**Figure 1a**).

**Fig. 1.**
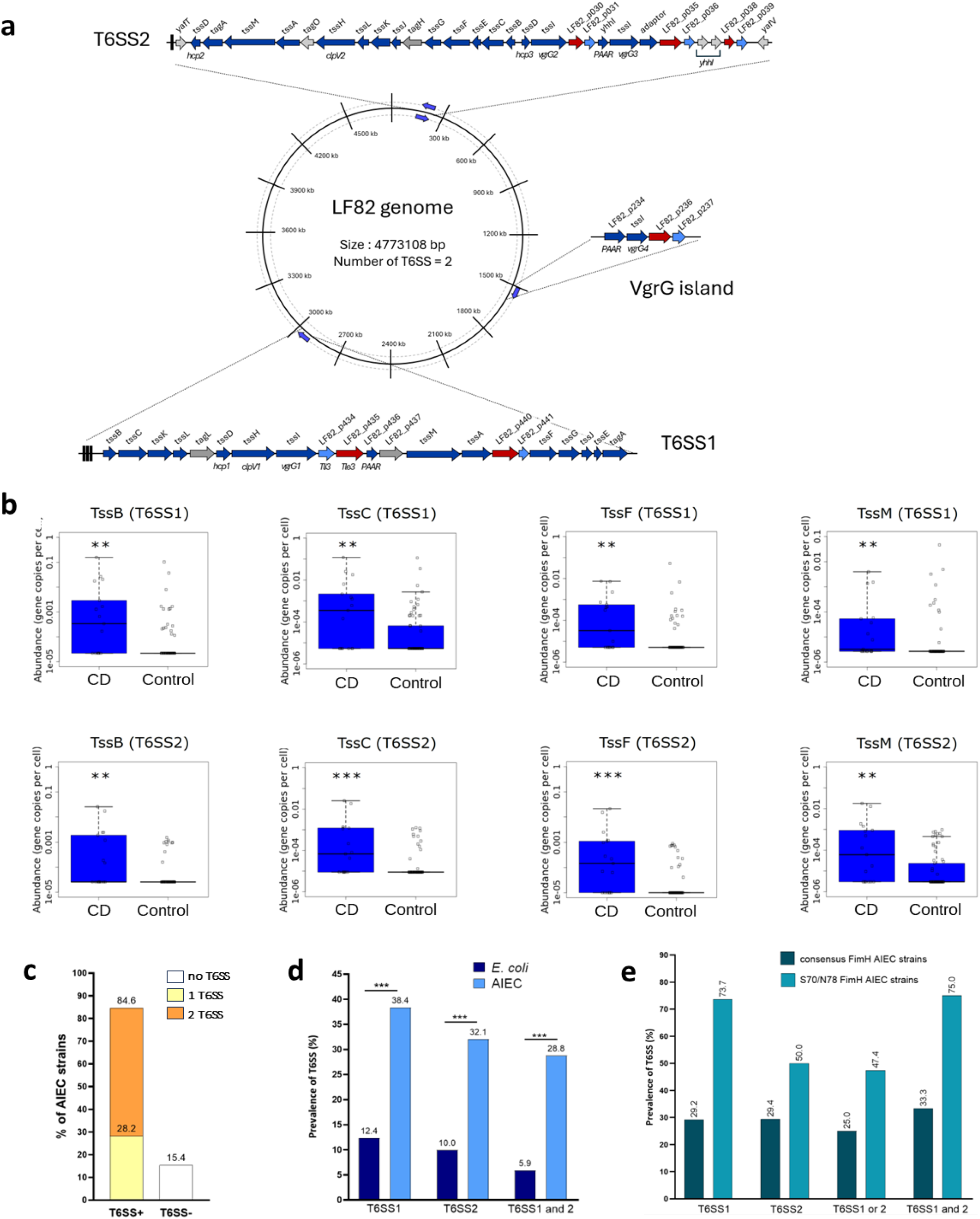
Prevalence of T6SS genes in CD patients’ microbiome compared to healthy controls. **a**. T6SS putative genes identified in LF82 genome. Genes encoding for core components of T6SS systems are in dark blue, genes identified as effectors are shown in red and associated immunity proteins are depicted in light blue. Two complete clusters of genes coding for T6SS are identified as T6SS1 and T6SS2 and some T6SS associated genes are found out of those clusters (VgrG island). **b**. Prevalence of T6SS genes in microbiota of CD patients (n=17) compared to healthy subjects (n=62). Meta-data was generated using MetaQuery webtool. ** *p*<0.01, *** *p*< 0.001 by Wilcoxon rank test. **c**. Prevalence of T6SS clusters in AIEC genomes (n=78) assessed by Macsyfinder TXSScan online tool. **d**. Prevalence of LF82-homolog T6SS clusters among *E. coli* sp. (n=1709) and AIEC pathovar (n=78). Genes from T6SS1 and T6SS2 systems of LF82 have been searched for in genomes taking into account the co-occurrence of four structural genes *tssB, tssC, tssF, tssG* with a minimum of 70% identity. Statistical significance was assessed by Chi square test (*** *p*<0.001, ns non significant). **e**. Prevalence of LF82-homolog T6SS clusters in AIEC strains (n=78) according to S70/N78 polymorphism on FimH.

The abundance of T6SS genes in CD patient’s intestinal microbiota was explored through the web-based server MetaQuery dedicated to quantitative analysis of specific genes in the human gut metagenome. Four structural genes essential for T6SS functioning were selected: *tssB* and *tssC* (structural proteins of the sheath), *tssF* and *tssM* (belonging to T6SS baseplate and membrane complex respectively). As two different copies of each gene were found in AIEC LF82 genome (**Figure 1a**), both sequences have been used for the analysis. MetaQuery generated the abundance of the eight selected sequences in metagenomes of 17 Crohn’s disease patients and in 62 healthy subjects. Each T6SS gene searched for was significantly enriched (p<0.01) in the microbiome of CD patients, indicating a probable higher prevalence of bacteria armed with T6SS in their microbiome and suggesting a potential role for these system in CD-associated symptoms and/or dysbiosis (**Figure 1b**).

### Prevalence of T6SS in AIEC strains isolated from ileal mucosa of CD patients

To better understand the importance of these systems in AIEC, a systematic research of T6SS clusters was first performed in genomes of 78 AIEC clinical isolates from our laboratory database using MacSyFinder TXSScan tool^21–23^. More than 85% of AIEC strains possess at least one T6SS cluster (**Figure 1c**). Then AIEC genomes and *E. coli* genomes available in the RefSeq database were screened for T6SS with significant similarity to those of LF82 reference strain. To identify complete gene clusters, strains were considered positive for T6SS homologous to those of LF82 when the sequence of all four genes *tssB, tssC, tssG* and *tssF* (respectively belonging to sheath, tube, and baseplate for the last two) were at a minimum of 70% identity with AIEC LF82 strain and when the maximal distance separating genes on genome sequence was less than 34 kb. According to these criteria, prevalence of at least one LF82-similar T6SS in *E. coli* RefSeq database is 17.2%, which is consistent with known data reporting that T6SS are found in around 25% of Gram-negative Pseumonadota^15^. Of note, co-occurrence of LF82-homologous T6SS1 and T6SS2 in RefSeq *E. coli* genomes is quite noticeable: among T6SS1-carrying *E. coli*, 47.6% also have a T6SS2 in their genome, and conversely 54.7% of T6SS2-carrying *E. coli* also have a T6SS1 in their genome. This could mean that a high proportion of *E. coli* often use or need both T6SS in their genomes. Among AIEC strains, 38.4% have a T6SS1 homologous to that of LF82 and 32.1% have a T6SS2 homologous to that of LF82. More than half of these strains (representing 28.8% of AIEC collection) carry both T6SS1 and T6SS2, versus 5.9% in *E. coli* RefSeq database (**Figure 1c**). This suggests that these two operons could play a role in AIEC phenotype.

Previous work from our laboratory reports that a significant proportion of AIEC clinical isolates display a pathoadaptative polymorphism on FimH gene encoding type 1 pili adhesin involved in bacterial attachment to their receptor at the surface of intestinal epithelial cells^34^. Therefore, we compared the prevalence of LF82-apparented T6SS1 and T6SS2 in AIEC strains with consensus FimH sequence versus those with S70/N78 mutation. Both T6SS1 and T6SS2 appear associated to pathoadaptation in FimH gene (**Figure 1e**), which could mean that T6SS might be associated with better AIEC adaption within host and higher virulence.

With a high percentage of AIEC possessing both LF82-apparented T6SS1 and T6SS2, the reference AIEC strain LF82 seems to be a good model to explore the role of those systems within gut microbiota.

Despite CD patients colonized by AIEC are at higher risk of post-operative recurrence^19^, clinical outcomes of patients from POP-Remind cohort (NCT03458195)^33^ were not associated with the presence or absence of LF82-homologous T6SS in the genome of their colonizing AIEC strain (**Figure S2**).

### Identification of AIEC LF82 T6SS expression conditions

In order for AIEC bacteria to take benefit from the T6SS in a complex ecological context, T6SS need to be expressed under gastrointestinal conditions. Hence, the TNO-gastro-intestinal model (TIM-1) was used to determine whether T6SS are expressed under intestinal conditions. Indeed, this computer-controlled *in vitro* model realistically simulates luminal conditions of successive gastro-intestinal compartments (stomach, duodenum, jejunum and ileum; **Figure 2a**). The chemical conditions of human digestion are replicated through the constant addition of different solutions mimicking organs secretions, and even physical conditions such as mixing, meal transit, removal of digested compounds and water, are imitated with different pumps and filters strategically placed in the model (**Figure 2a**).

**Fig. 2.**
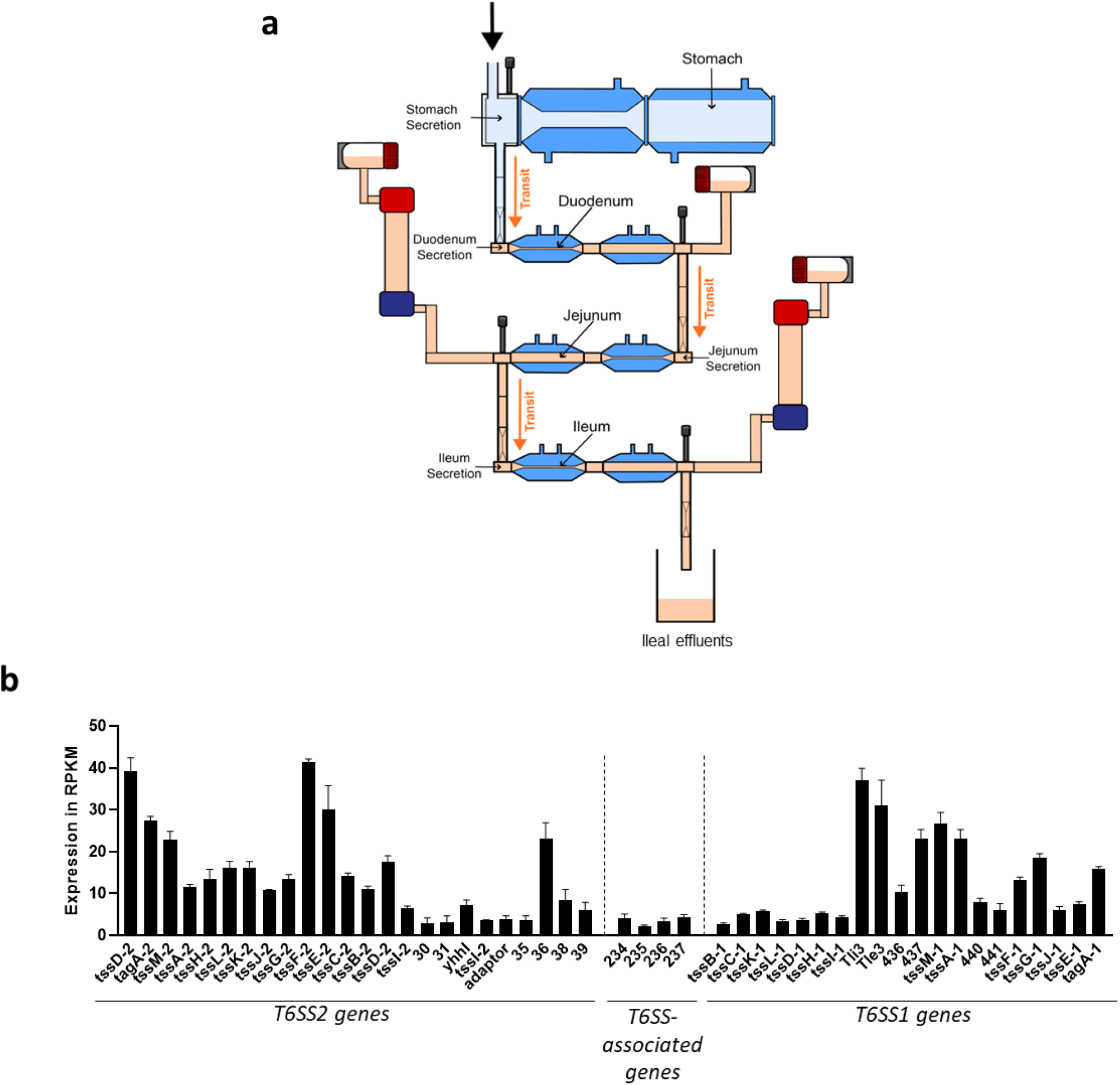
LF82 T6SS expression conditions. **a**. Schematic representation of the main compartments of the *in vitro* upper tract digestive model TIM-1. LF82 inoculation is represented with the black arrow. Five hours post-inoculation, ileal effluents were used for transcriptomic analysis. **b**. Expression of T6SS genes measured in TIM-1 at 5 hours post inoculation in RPKM unit. Results are mean ± SEM of three independent experiments.

Considering that AIEC are most often isolated from patients’ ileum and that this system is very complete for replicating human small intestine conditions, we introduced AIEC LF82 WT strain into the stomach of the TIM-1, and 5 hours later ileal effluents were collected to perform Illumina RNA sequencing. Results showed that genes within the same cluster are not equally expressed at T=5h. For T6SS2, structural proteins-coding genes (*tssD-2* to *tssI-2*) are expressed, meaning that LF82 can probably assemble a full T6SS2. Additionally, for T6SS1 cluster, expression of genes is higher starting from p434-encoding gene, suggesting the presence of an intra-cluster regulation, yet not identified (**Figure 2b**). However, both systems are clearly expressed in intestinal conditions, therefore suggesting that LF82 can potentially utilize them in the digestive tract.

To better understand the conditions under which the two T6SS-encoding operons are expressed, a promoter assay was performed. Three promoters have been identified, the first one for T6SS1 operon upstream of *tssB-1* gene and two other ones for T6SS2 operon between *tssB-2* and *tssD-2* genes (**Figure 3a**). Promoter regions have been cloned in pRS550 reporter plasmid and transformed into AIEC LF82 WT strain in order to evaluate promoter activity in various growth conditions, chosen to mimick AIEC lifestyle in intestinal (variation of pH, high osmolarity, chelation of metal ions, presence of sodium cholate only or presence of bile salts (i.e. sodium cholate and deoxycholate mixture), presence of reactive oxygen species, disruption of bacterial membrane) and intracellular contexts (i.e. acidic pH, mimic vacuole, reactive oxygen species). Supplementation with 2% of bile salts was the only condition leading to elevated β-galactosidase activity, especially for pRS-TS1 and pRS-TS2left. For pRS-TS2right, the level of induction is the lowest between the three promoters tested but still significantly higher than the control group (**Figure 3b**). Bile salts, therefore, appear to play a key role in T6SS promoter induction and most likely in the transcription of T6SS-encoding operons.

**Fig. 3.**
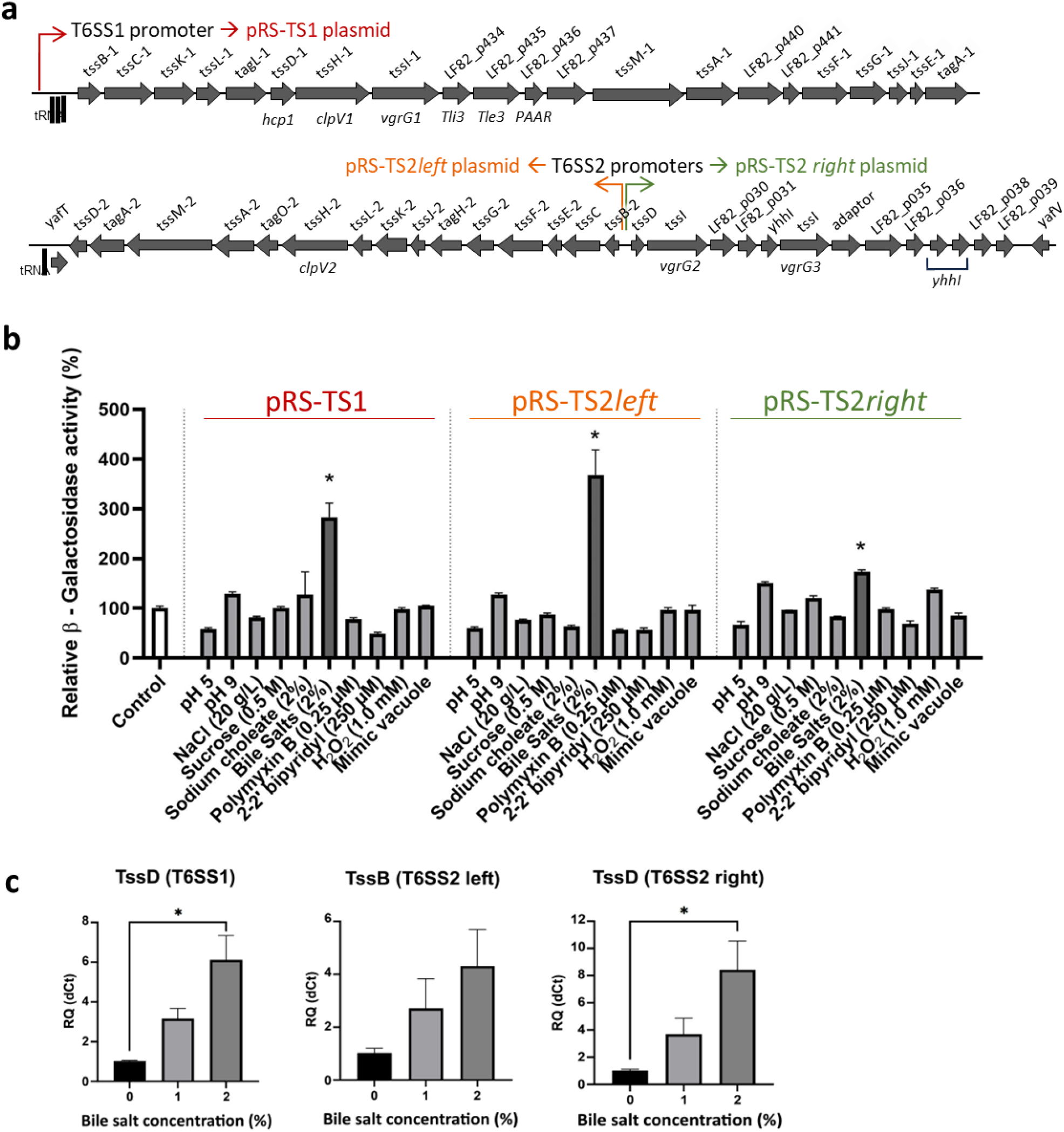
T6SS LF82 promoter induction conditions. **a**. Three promoters have been identified in T6SS LF82 clusters as shown on the illustration. **b**. Promoter reporter assay. Promoter regions were cloned into a pRS550 plasmid and transformed into LF82 WT strain. Promoter induction is measured in various conditions through % of beta-galactosidase activity and compared to LF82 WT strain carrying an empty pRS550. Results are mean ± SEM of 3 replicates and statistical significance was assessed by one-way ANOVA followed by Dunnett’s multiple comparison test (* p<0.05). **c**. RT-qPCR analysis for expression of T6SS structural protein-coding genes after incubation in the presence of various bile salt concentrations; *idnT* was used as housekeeping gene. Results are perecntages of ΔCt relative quantification (mean ± SEM of from 3 independent experiments) and statistical significance was assessed by Kruskal-Wallis test (* p<0.05).

To confirm bile salts involvement in T6SS gene expression, RT-qPCR was performed on LF82 incubated with increasing bile salts concentrations. Of note, more than 2% bile salts were deleterious and caused death of a significant fraction of bacteria (data not shown). Genes tested were *tssB-2, tssD-1* and *tssD-2* encoding for proteins of the sheath and tube of T6SS, thus they are essential for T6SS assembly and firing. For the three genes tested, expression was increased by bile salts in a dose-dependent manner (**Figure 3c** and **S3**), confirming that bile salts represent a major factor in regulation of LF82 T6SS gene expression. Since T6SS are expressed in intestinal conditions, they could be a valuable asset for AIEC to compete with resident gut microbiota.

### Impact of LF82 T6SS on mice intestinal microbiota

To evaluate the impact of AIEC T6SS on intestinal microbiota, we administered LF82 WT strains and its isogenic T6SS mutants to healthy mice without antibiotic treatment (usually administered to disrupt colonization resistance). C57BL/6N mice were daily orally infected with LF82 WT strain, LF82 ΔΔT6SS1.2 (isogenic mutant deleted for both T6SS clusters) or LF82 Δ435 deleted for the *Tle3* effector gene (**Figure 4a**). Illumina-based sequencing of the V3-V4 region of the 16S rRNA was next performed on feces of infected mice. At day 0, no clusterization was observed between the three groups of mice, showing the absence of cage effect (**Figure S4**). At day 10, LF82 WT-infected mice clustered differentially compared with mice infected with LF82 ΔΔT6SS1.2 mutant (permanova q-value = 0.0465) or LF82 Δ435 mutant (permanova q-value = 0.0465), suggesting that the presence of T6SS in LF82 could modulate the intestinal microbiota composition (**Figure 4b**). There was no significant clusterization between LF82 ΔΔT6SS1.2 and LF82 Δ435 mutants (permanova q-value = 0.1). No significant alteration of alpha diversity was detected between day 0 and day 10 (Chao1, Shannon and Simpson indexes, data not shown). Although there was no major disruption in microbiota composition, specific changes were nonetheless identified in the microbiota. Among all the bacterial groups identified, very few were equally distributed among the three experimental conditions and the abundances of several groups were significantly shifted between conditions. *Staphylococcus* and especially *Staphylococcus sciuri* were significantly more abundant following infection with AIEC LF82ΔΔT6SS1.2 mutant than with the WT parental strain (**Figure 4c**), suggesting that they could be direct or indirect targets of LF82 T6SS. However, *Staphylococcus* do not seem to be targeted by p435 effector. Conversely, *Lactobacillus* genera appeared to indirectly benefit from the presence of LF82 T6SS, possibly through T6SS effects on other microbiota members (**Figure 4c**). Although inter-individual heterogeneity limits statistical significance in our experiment, non-significant trends of interest can also be detected : *Bilophila, Oscillospira, Coprococcus, Dorea* and to a lesser extent *Roseburia* and *Mucispirillum schaedleri* tended to be more abundant in mice infected with LF82ΔΔT6SS1.2 or LF82 Δ435 mutants compared to those infected with LF82 WT, although this was not significant (**Figure S5**). In mice infected with single T6SS mutants LF82 ΔT6SS1 or LF82 ΔT6SS2, no significant difference of abundances was noticed, except for *Staphylococcus* whose abundance was significantly increased upon infection with LF82 ΔT6SS2 (**Figure S6**). Altogether, these results suggest that LF82 T6SS play a role in modulating the microbial ecosystem upon LF82 infection.

**Fig. 4.**
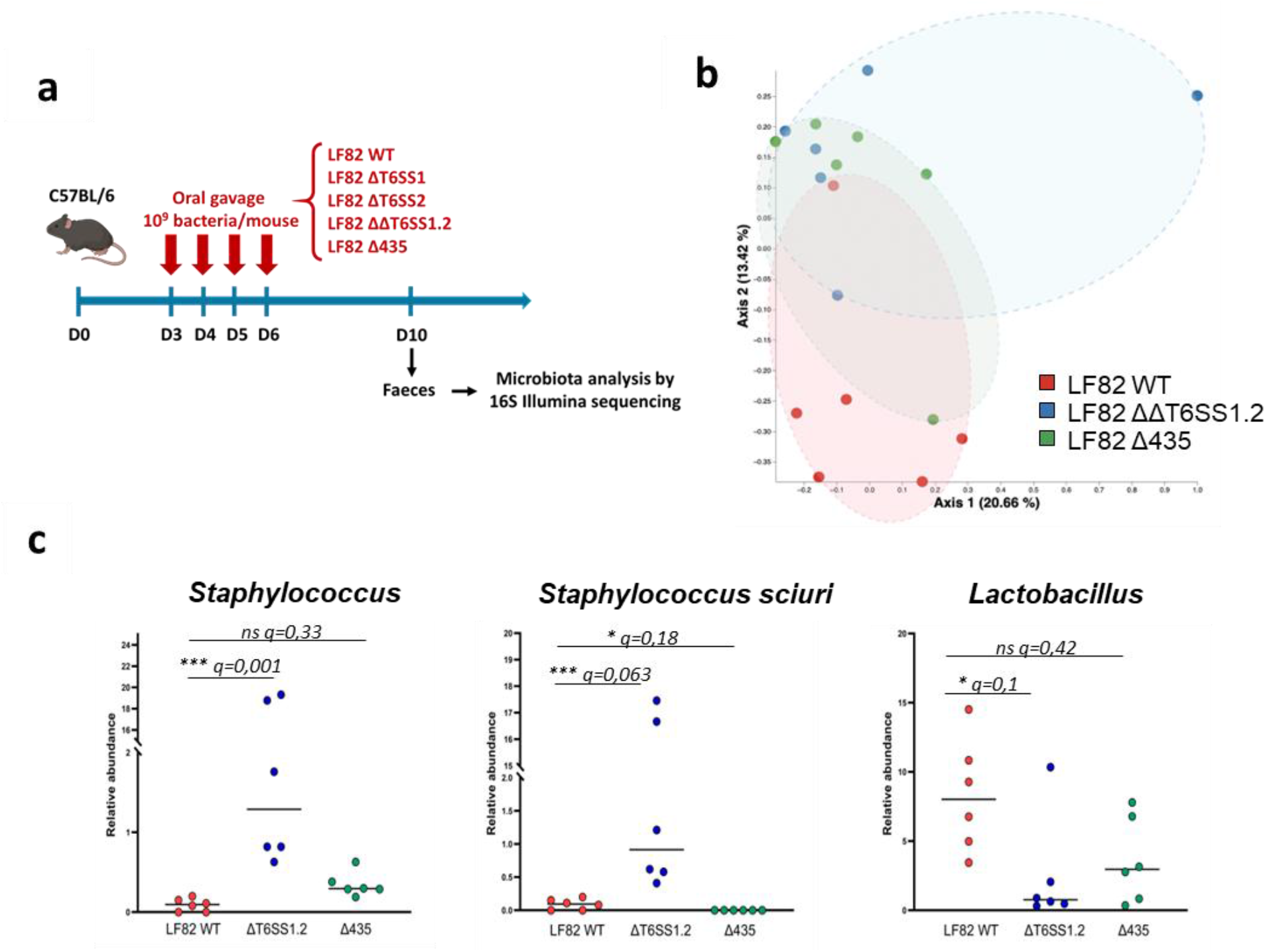
Effect of T6SS from LF82 on mouse microbiota composition. **a**. Design of *in vivo* experiment. C57/BL6 mice were gavaged for four consecutive days with LF82 WT, LF82 ΔT6SS1.2, LF82 ΔT6SS1, LF82 ΔT6SS2 or LF82 Δ435 strains. Faeces samples were collected and used for 16S rRNA Illumina sequencing. Microbiota alpha and beta-diversity were analysed. **b**. PCoA analysis of microbiota beta-diversity in fecal samples on day 10 (D10). Dots indicate mice infected with LF82 WT strain (in red), with LF82 ΔT6SS1.2 (in blue) or with LF82 Δp435 (in green). **c**. Relative abundance of bacterial groups significantly shifted in mice infected with LF82 ΔT6SS1.2 and/or LF82 Δ435 compared to those infected with LF82 WT; ns: non-significant, q-values are indicated.

## Discussion

The aim of the study was to explore the role of T6SS, a widespread virulence factor in Gram negative bacteria. The important prevalence of T6SS clusters in AIEC suggests that these secretion systems are not only related to *E*.*coli* lifestyle but that T6SS could rather play a role in AIEC virulence, possibly by conferring an advantage for competition with host microbiota, contributing to AIEC encroachment within the intestinal ecosystem or driving CD-associated dysbiosis. This hypothesis is reinforced by the co-occurrence of LF82-homolog T6SS with pathoadaptive mutation on FimH virulence factor. Therefore, we first sought to determine whether T6SS1 and T6SS2 of LF82 AIEC reference strain were expressed under intestinal conditions and then if this expression impacted gut microbiota diversity and composition *in vivo*.

It has been observed in the literature that some bacteria constitutively express their T6SS, whereas others maintain strict regulation of T6SS genes, which seems to be the case for LF82^35^. Testing expression of T6SS operons in intestinal conditions was necessary to verify that those systems were supposably active in this context, and we identified bile salts as a major factor driving T6SS expression. All bacteria within intestinal microbiota will be in contact with bile salts and respond differently to this environmental factor. Bile salts may serve as signals that identify intestinal environment or induce a stress on some intestinal bacteria^36,37^. As a matter of fact, bile salts drive the expression of several bacterial loci and can either promote or repress expression of bacterial virulence genes. For example, a fraction of bile salts, deoxycholate, represses invasion-related genes in *Salmonella*^38^ whereas it promotes virulence genes expression in *Campylobacter jejuni*^39^. Regarding specifically T6SS, bile salts have been reported to regulate their expression or activity in enteric pathogens such as *Salmonella* Typhimurium^40^ and *Vibrio cholerae*^41^. The underlying mechanisms are not elucidated, although transduction pathways have already been found responsible for bile salts-mediated regulation of other virulence genes, notably VtrA/VtrC co-component periplasmic complex promoting T3SS expression in *Vibrio parahemolyticus*^36,42^. The molecular basis of T6SS operons regulation in LF82 strain remain to be further explored, especially as modulation of the global regulator ppGpp has recently been reported to govern LF82 T6SS expression ^43^. Of note, our observations suggest the presence of internal promoter/regulatory sequences, as the expression levels measured are unequal along the operons. This would allow other intestinal factors (mucins, pH, oxygen availability and reactive oxygen species…) to participate in gene expression regulation. Interestingly, several genes encoding immunity proteins (p36, p434, p441) seem highly expressed compared to their neighbours. Additionally, such environmental factors could also modulate T6SS activity without affecting gene expression. As a result, T6SS-mediated virulence of AIEC bacteria would be finely tuned according to the location of the bacteria inside the gastro-intestinal tract.

Our results clearly show the impact of LF82 T6SS on microbiota diversity and composition, although we did not observe a drastic dysbiosis as described for example in patients or in mice submitted to colitis. This kind of T6SS-mediated modulation, at species or genus level only, is not surprising given the short time-course of the experiment and the lack of antibiotic-preconditioning of mice. This has already been reported for example with *Klebsiella pneumoniae*^44^. When considering our whole results, some variations are consistent between conditions and could be related to one T6SS in particular: abundance of *Staphylococcus* is significantly increased upon infection with LF82 ΔΔT6SS1.2 double mutant as well as with LF82 ΔT6SS2, but not with LF82 ΔT6SS1 nor with LF82 Δp435, suggesting that *Staphylococcus* could be a target of LF82 T6SS2 though secretion of antibacterial effectors that remain to be characterized. This is unexpected as T6SS are reported to target most often Gram-negative bacteria, although in the last years T6SS of *Pseudomonas chlororaphis* and *Acinetobacter baumannii* have been reported to kill Gram positive preys^45,46^. Of note, T6SS can also act through exploitative competition by secreting effectors promoting uptake of rare metallic ions and enhancing bacterial fitness^47^, and we cannot exclude this kind of process in our experiment. It is also possible that a large part of our observations is related to indirect impact of T6SS, indeed the expansion of target microorganisms in a context of infection with T6SS-deleted LF82 could in turn disadvantage their competitors; or conversely T6SS-mediated elimination of some bacteria could free a niche not only for T6SS-harboring AIEC but also for other members of microbiota. Moreover, in some case T6SS can target both procaryotic and eucaryotic cells by secreting either several effectors with distinct targets or transkingdom effectors as reported in *Pseudomonas aeruginosa*^9^. Some effectors of LF82 T6SS probably act on host cells, which may modulate also the inflammatory status of gut mucosa and in turn influence resident microbiota.

Finally, it is interesting to note that among microorganisms showing a non-significant trend to decrease in the presence of LF82 T6SS, several are known to be short-chain fatty acids-producers with anti-inflammatory properties and to be under-represented in CD patients’ microbiota: *Roseburia, Oscillospira, Coprococcus, Dorea*^48^. In contrast, hydrogen sulfide-producing *Bilophila* and mucus-penetrating *Mucispirillum* are usually considered as pro-inflammatory and potentially harmful bacteria^49,50^. Therefore, AIEC T6SS modify the balance between beneficial bacteria and pathobionts within gut microbiota. However, this is probably not the only function of T6SS in LF82 virulence, since previously published data suggest that LF82 T6SS are also expressed intracellularly in macrophages^51^. Except for p435 (Tle3), the T6SS effectors of LF82 are not characterized yet, asking the question of their possible impact towards host cells during mucosal colonization.

In this study, we investigated the role of T6SS in the virulence of AIEC isolated from Crohn’s disease patients and demonstrated that T6SS genes are expressed in gastro-intestinal conditions (bile salts) and can impact directly or indirectly intestinal microbiota composition, while the exact role on LF82 pathogenesis remains to be identified. Before concluding, it has to be noted that 40% of the strains in our AIEC collection possess T6SS systems that share less than 70% homology to those of LF82 but which may still be involved in AIEC phenotype. Extended understanding of the pathogenicity and physiology of AIEC could undoubtedly aid in the development of novel drugs targeting T6SS as a promising new strategy for controlling gut colonization and AIEC-induced inflammation in CD patients.

## Acknowledgments

The authors thank the Genom’IC platforms (Inserm U1016, Paris, France) for their help in library sequencing, and all the investigators from the Remind Group. Parts of this work have been presented as posters in the 16^th^ national congress of the French Microbiology Society (2021, Nantes, France) and in Cell Symposia congress « Infection biology in the age of the microbiome » (2023, Paris, France).

This work was supported by Inserm, INRAe, Clermont-Auvergne University and by grants from the French National Research Agency (ANR-18-CE15-0013 EffAIECts, ANR-21-CE15-0042-01 EMULBIONT, ANR-22-CE17-0054 TOPIC). BC’s laboratory is supported by a Starting Grant (Grant Agreement Invaders No. ERC-2018-StG-804135) and a Consolidator Grant (Grant Agreement InterBiome No. ERC-2024-CoG-101170920) from the European Research Council (ERC) under the European Union’s Horizon 2020 research and innovation program, ANR grants EMULBIONT (ANR-21-CE15-0042-01) and DREAM (ANR-20-PAMR-0002), DIM BioConvergence for Health (BioConvS) and Région Ile-de-France and the national program “Microbiote” from Inserm. No funders had any role in the design of the study and data collection, analysis and interpretation, nor in manuscript writing. REMIND study has been supported by grants from MSD France, Association François Aupetit and the Helmsley Charitable Trust.

## Author contributions

NB, EB and LJ obtained the funding and supervised the project. MA, PS, ABu and Remind study Group investigators enrolled and followed-up patients. NB, EB, LJ and BC designed the experiments. HB, SM, GS, CC, MR and SD conducted the experiments, collected and analyzed the data. EP and IW processed the RNAseq data. BC analyzed the 16S sequencing data. RB and ABi analyzed genome databases for T6SS prevalence. BP performed statistical analysis on clinical data. HB and EB wrote the article. LJ, BC, SD and JD critically revised and edited the manuscript. All authors read and approved the final manuscript.

## Data availability statement

16S sequencing data and datasets corresponding to the main figures of this paper are available upon request (https://doi.org/10.57745/WWGSAS).

