## Supplemental Figures S1 to S6 for "Type 6 secretion systems of adherent-invasive *Escherichia coli* isolated from Crohn’s disease patients shape intestinal microbiota"

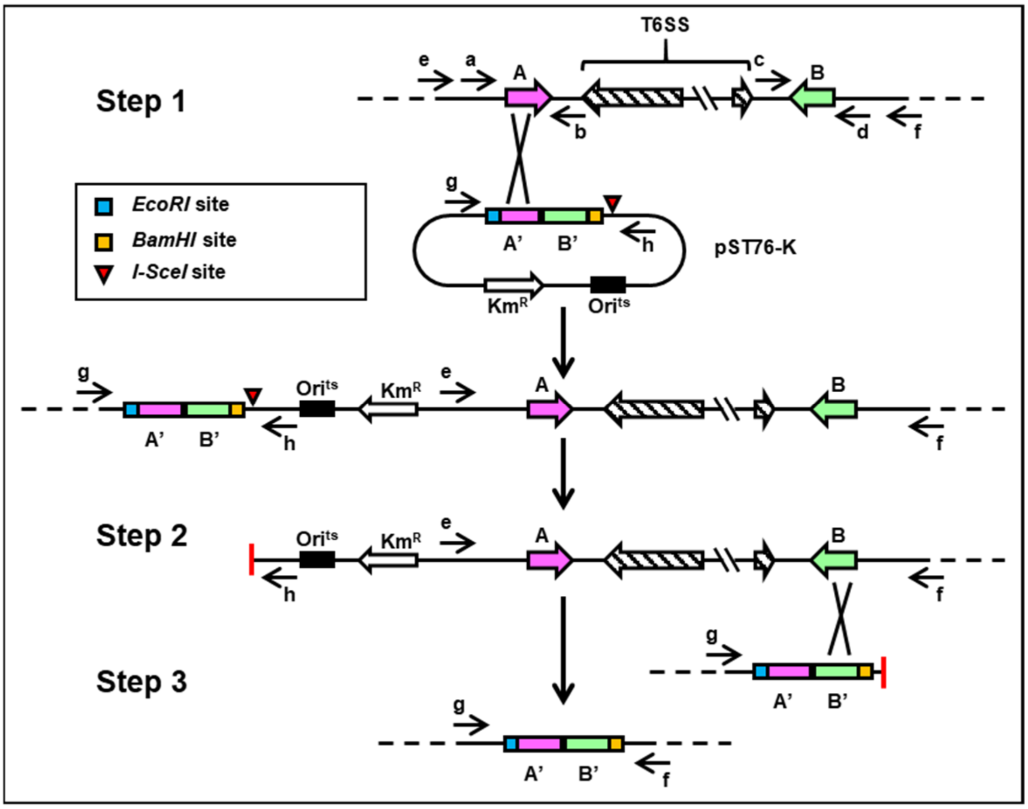


**Figure S1: Strategy for T6SS cluster deletion in LF82 strain**

**Step 1.** Cloning of upstream (A’) and downstream (B’) regions that frame the T6SS targeted into the mutagenesis suicide pST76-K plasmid carrying *I-SceI* recognition site. The mutagenesis plasmid (pST76-K-TS1 or TS2) is introduced into wild-type *E. coli* LF82 and integrated into upstream region (A) by homologous recombination. Integration can also occur to downstream or both region (not shown). **Step 2.** Plasmid pDAI-SceI, which expresses the *I-SceI* nuclease (not shown) is introduced into the single cross-over insertion strain. *I-SceI* expression results in cleavage at the *I-SceI* recognition site, producing a double strand break in the chromosome. This DNA break stimulates the host recombination/repair machinery causing a second recombination event between B and B’ (desired deletion) or between A and A’ (back to wild type strain). In any type of repair, the integrated plasmid is lost along with the kanamycin gene. **Step 3.** PCR amplification performed on recovered colonies that are kanamycin sensitive confirm that the desired deletion of T6SS1 or T6SS2 has been obtained. a, b, c, d, e, f, g and h represent PCR primers.

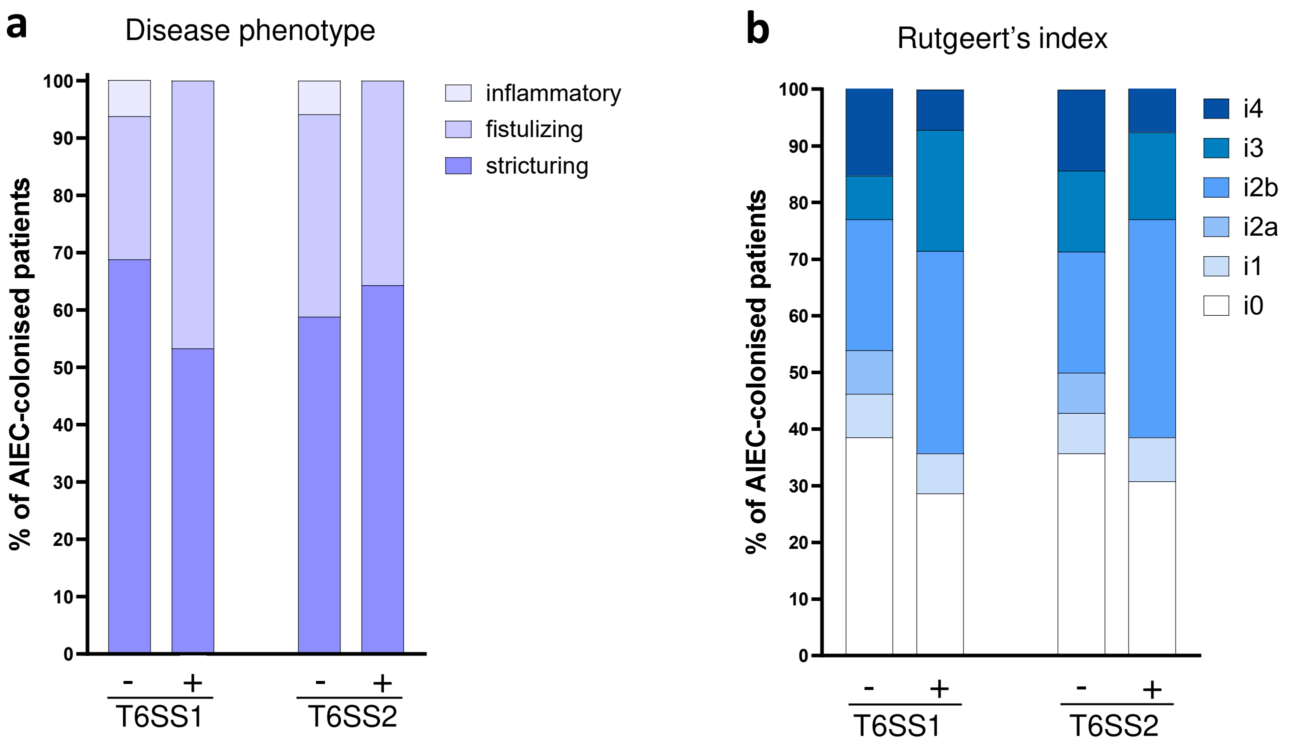


**Figure S2 : Clinical outcomes of CD patients according to the presence or absence of T6SS homologous to those of LF82 strain.**

CD patients from POP-Remind cohort undergo ileal resection with prospective follow-up. Patients with AIEC ileal colonization were identified by isolating *E. coli* from surgical ileal samples and assessing adherent-invasive phenotype in vitro[19]. Clinical outcomes of these patients were analyzed according to the presence or not of LF82-homologous T6SS in the genome of their colonizing AIEC strain.

a. Disease phenotype was evaluated at enrollment.

b. Endoscopic examination was performed at 6 months following surgery to grade endoscopic post-operative recurrence according to the Rutgeerts' index : i0 = no lesion in the distal ileum ; i1 = less than 5 aphthous lesions in the distal ileum ; i2a = ulcers up to 1 cm confined to ileocolonic anastomosis ; i2b = more than 5 aphthous lesions with normal mucosa between the lesions, or skip areas of larger lesions ; i3 = diffuse aphthous ileitis with diffusely inflamed mucosa between the multiple aphthae ; and i4 = diffuse inflammation, with large ulcers and/or nodules/cobble and/or narrowing/stenosis[50].

No statistical association was detected by $\chi^2$ or Fisher's exact tests.

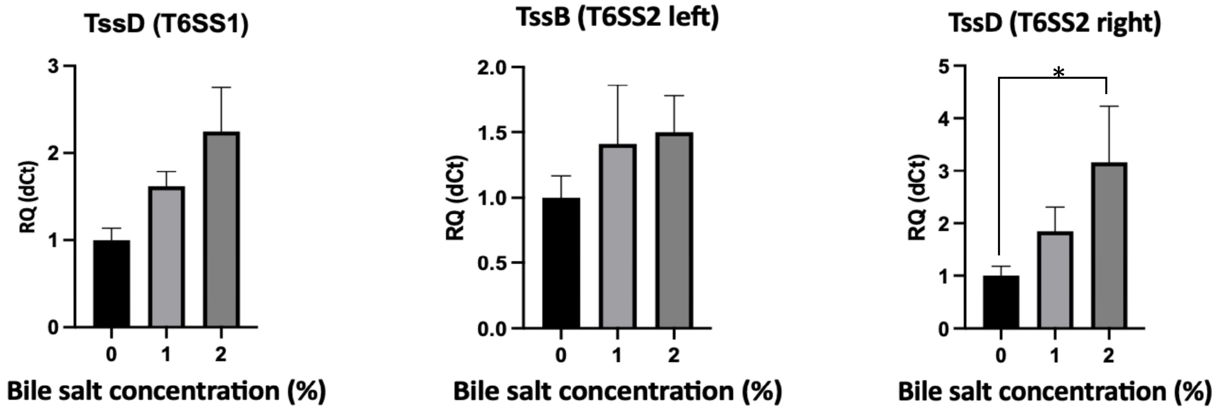


**Figure S3: Expression of T6SS genes in the presence of bile salts**

RT-qPCR analysis for expression of T6SS structural protein-coding genes after incubation in the presence of various bile salt concentrations; *cysG* was used as housekeeping gene. Results are percentages of ΔCt relative quantification (mean ± SEM of from 3 experiments) and p-values were determined by Kruskal-Wallis test (* $p<0.05$).

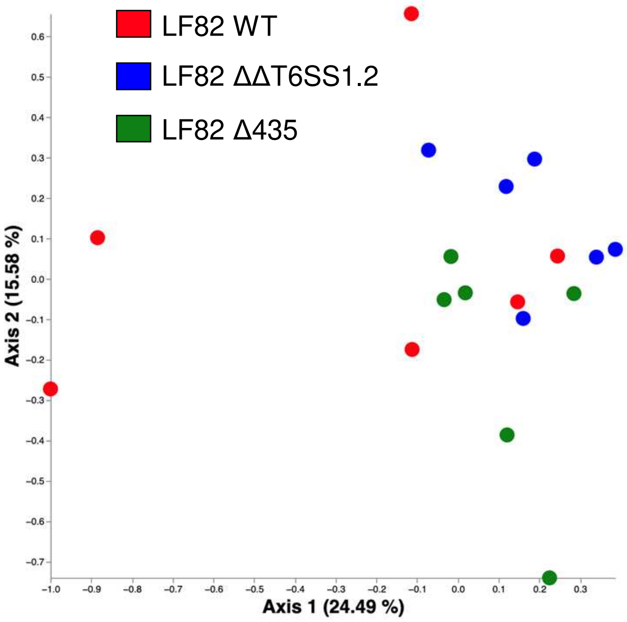


**Figure S4: PCoA analysis of microbiota beta-diversity at day 0 in fecal samples of mice**
LF82 WT, ΔT6SS1.2 and LF82 Δ435 groups of mice are shown, no significant clusterization is visible.

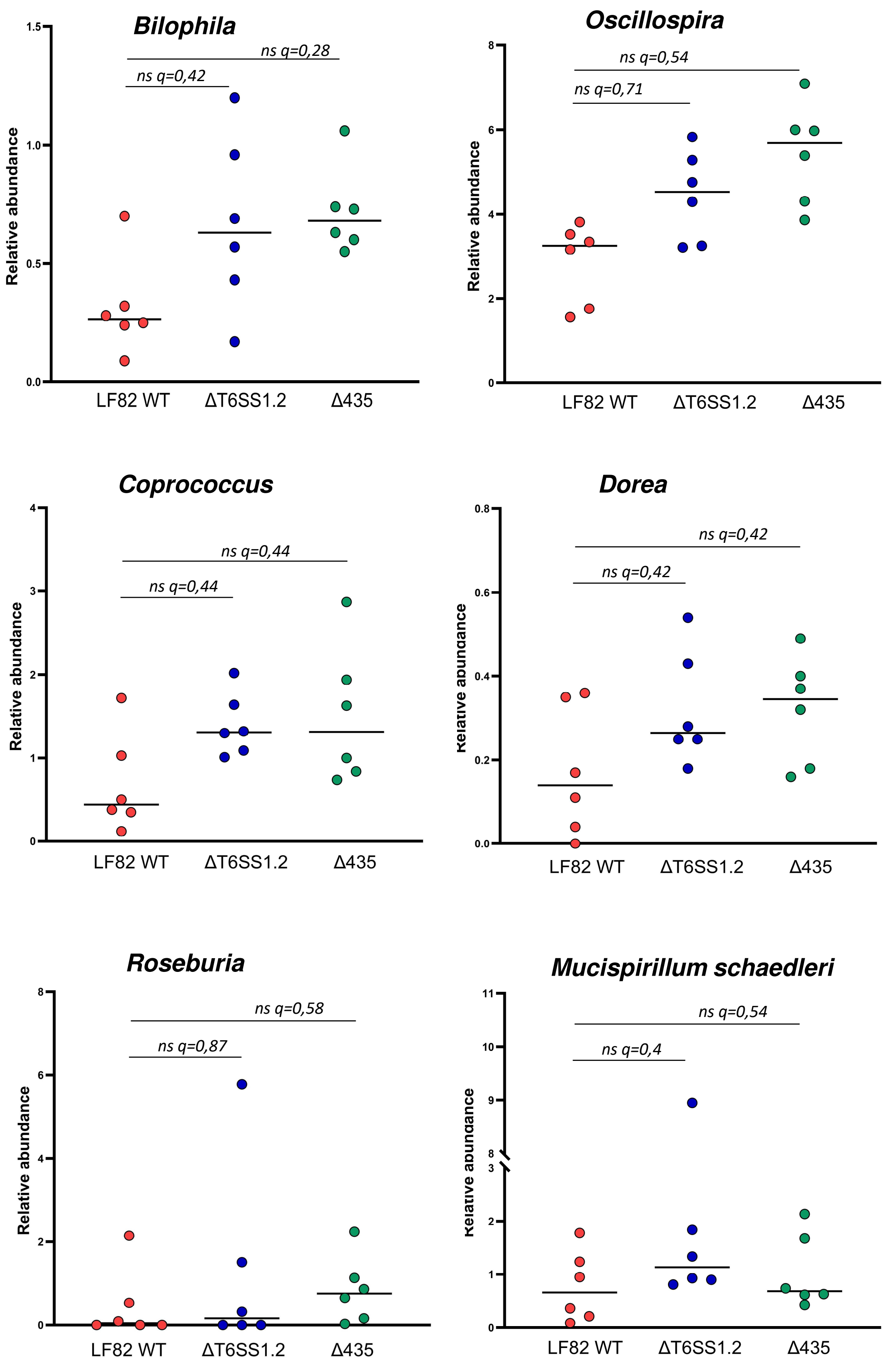


**Figure S5: Relative abundances of bacterial genera showing a tendency to shift in mice infected with LF82 ΔT6SS1.2 and/or LF82 Δ435 compared to those infected with LF82 WT.** ns: non-significant, q-values are indicated.

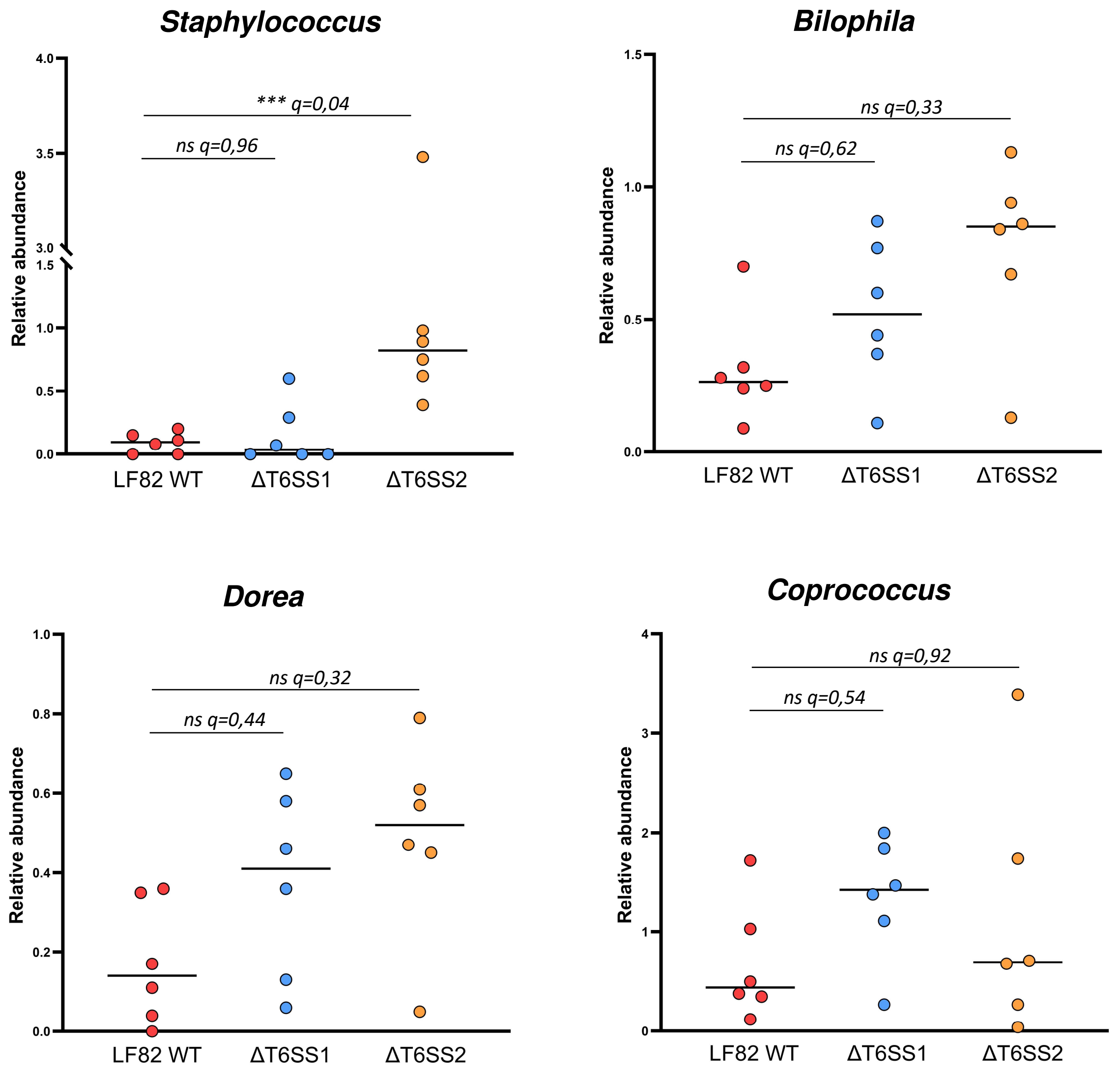


**Figure S6: Relative abundances of bacterial genera of interest in mice infected with LF82 ΔT6SS1 or LF82 ΔT6SS2 compared to those infected with LF82 WT.**
ns: non-significant, q-values are indicated.
